# Three-dimensional tissue strain measurement using a row-column array during biaxial testing of excised ventricular myocardium

**DOI:** 10.1101/2024.11.20.624360

**Authors:** Xavier Navy, Zhiyu Sheng, Kang Kim, John M. Cormack

## Abstract

**Objective:** To implement and validate a 3D volume imaging sequence and 3D strain estimation procedure for enhanced biaxial mechanical testing of excised ventricular myocardium.

**Methods:** One right- and one left-ventricular excised porcine myocardium specimens were tested using dual-loading protocol quasi-static biaxial mechanical testing. During biaxial testing, volume ultrasound images were acquired using a row-column addressed array probe using a synthetic aperture imaging sequence. Volume ultrasound images were used to compute tissue deformation using 3D correlation-based ultrasound speckle tracking. Ultrasound-derived tissue strains were validated against repeated measurements using typical standard optical camera imaging of the surface deformation of the specimen.

**Results:** Synthetic aperture imaging with the RCA yielded artifact-free and uniform ultrasound speckle for high-fidely speckle tracking of tissue deformation. Ultrasound-derived tissue strain is in good agreement with ground-truth camera-derived surface strain measurements (root-mean-square error is 1.6% strain).

**Conclusion:** High-fidelity 3D strain measurement with ultrasound imaging is accurate and can enhance biomechanical insights from biaxial experimentation, especially in large tissues such as porcine and human myocardium where assumptions of plane stress and incompressibility may not apply.

## 1. Introduction

Biaxial mechanical testing is a popular benchtop experimental technique for determining anisotropic mechanical properties of excised soft tissues,^1^ including ventricular myocardium,^2,3^ bladder,^4^ and vascular^5^ applications. In current practice, high fidelity force measurements are coupled with tissue deformation measurements by the tracking of superficial markers in optical camera images. Thus the measurement technique is not sensitive to out-of-plane motion and deformation, and the inverse problem of determining tissue mechanical properties must be made tractable by employing assumptions such as plane stress and incompressibility that may not apply in all cases.^6,7,8^

Our group has demonstrated coupling ultrasound (US) imaging with biaxial mechanical testing of excised myocardium, wherein the 3D displacement and strain were obtained by linear^9^ or rotational^8^ scans of linear US imaging arrays along with three-dimensional US speckle tracking (3DUST) techniques.^10^ Mechanical scanning of the US imaging array resulted in long data acquisition times, thereby limiting testing to quasistatic deformations and limiting the number of deformed configurations that could be produced during a reasonable testing duration.

In this letter we introduce and validate 3D strain measurement during biaxial testing of ventricular myocardium without mechanical scanning by acquiring volume 3D US images using a row-column addressed imaging array (RCA) and synthetic aperture beamforming.^11^ 3DUST is used to obtain displacement and strain throughout the specimen volume. Strain calculated by 3D US imaging is validated against strain measured on the surface using conventional camera imaging.

## 2. Methods

### 2.1 Tissue preparation and biaxial testing

The right and left ventricular free walls of a fresh frozen porcine heart were removed from the septum, and 20 *×* 20 mm^2^ square sections were excised from near the tricuspid and mitral valves, respectively.^8^ The right ventricular (RV) sample was left as full thickness (approximately 8 mm) for testing, while the left ventricular (LV) sample was thinned to approximately 5 mm thickness including the epicardium.

Each tissue sample was loaded into the biaxial testing machine (BioTester 5000, CellScale, Ontario, Canada) using fishhooks and pulleys to reduce shear loading (Fig. lA).^1,6^ The sample was submerged in de-ionized water at approximately room temperature. Two-protocol quasistatic biaxial stretching was employed^8,9^ in which the actuator displacements in the two loading protocols have ratios of l:l and l:2 (X:Y), respectively. Preconditioning and re-tensioning procedures^8^ were followed prior to the testing cycles to account for tissue damage that occurred during preconditioning. Three stretched configurations were obtained for each of the two stretching protocols (6 stretched configurations total per sample).

### 2.2 Ultrasound volume imaging and 3D strain calculation

An RCA (RC6gV, Vermon SA, Tours, France, 6.0 MHz center frequency, 256 channels, 25.4 *×* 25.4 mm^2^ footprint) was poised above the tissue prior to biaxial testing (Fig. 1B) with lateral and elevational axes aligned with the BioTester’s *X* and *Y* actuators. Ultrasound imaging during biaxial testing was controlled by a programmable research ultrasound imaging platform (Vantage 256, Verasonics, Kirkland, WA, USA).

Volume images were acquired in the reference configuration (after preconditioning and re-tensioning) and while the sample was held stretched in each of the stretched configurations (7 total volume images per sample tested). Imaging was performed with a synthetic aperture (SA) approach.^11^ The imaging sequence is to transmit from each individual element sequentially. Immediately after each row (column) element transmission, all of the column (row) elements record the echo signals simultaneously. Delay-and-sum (DAS) beamforming is used to reconstruct the ultrasound volume image.

The ultrasound image signal used for 3DUST was B mode brightness (in decibel units). 3DUST was performed in 13 *X*-*Z* planes between *Y* = −9 mm and *Y* = 9 mm in steps of l.5 mm.^8,10^ The cubic kernel for 3DUST had side length of 3 mm, and a search distance of 1.5 mm in each direction was employed. Tissue motion in three-dimensions throughout the tissue volume was determined in this way between each deformed configuration of each stretch protocol.

The trajectory **x**(**X**) of a particle that began at **X** = [*X, Y, Z*]^*T*^ was obtained by accumulating displacements measured from 3DUST. The positions **x**_*i*_ of particles in configuration *i* were found iteratively from

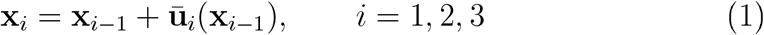

where **ū**_*i*_(**x**_*i*−1_) are the displacements measured by 3DUST between deformed configurations and **x**_0_ = **X**. Interpolation is used to estimate **ū**_*i*_(**x**_*i*−1_) for calculations if the deformed position **x**_*i*−1_ does not fall onto the grid on which 3DUST was performed. The total displacement throughout a deformation protocol is then **u**_*i*_(**X**) = **x**_*i*_(**X**) − **X**. Displacement accumulation was performed for tissue particles that have 3DUST normalized correlation coefficients greater than 0.35 and B mode brightness greater than −50 dB, respectively, throughout the loading protocol.

The displacement gradient **H** = *∂***u***/∂***X** was estimated by fitting a linear model to the total displacement **u**(**X**) in each stretched configuration obtained from accumulation. The region of fit was a rectangular parallelepiped volume with l4 mm on each lateral (*X*) and elevational (*Y*) sides, and included the entire tissue sample thickness in the axial (*Z*) direction as identified in the ultrasound volume image of the reference configuration. The coefficients of the linear models fit to each displacement component are the elements of **H**:

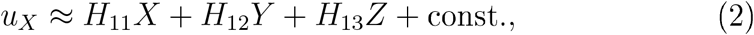

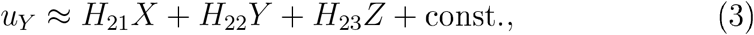

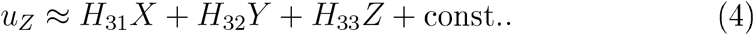

The deformation gradient tensor was calculated as **F** = **I** + **H**, where **I** is the identity tensor. The finite strain tensor **E** was obtained from the displacement gradient using

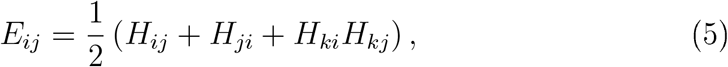

where the index summation convention is used.

### 2.3 Validation of strain calculation

After biaxial stretching while imaging with the RCA, the RCA was removed and the optical camera put in place aimed at the center of the tissue surface (Fig. lC).^3,8,9^ The test cycles were repeated while the camera captured images at a rate of 1 Hz. Displacement of the tissue surface in two dimensions was estimated using the Labjoy software (CellScale),^8^ and inplane displacement gradient and strain computed using Eqs. (2)-(5).

### 2.4 Stress calculation

The second Piola-Kirchhoff (P-K) stress **S** = **PF**^−1^ where **P** is the first P-K stress, is obtained by assuming plane stress and using the formulation given by Sommer et al.:^6^

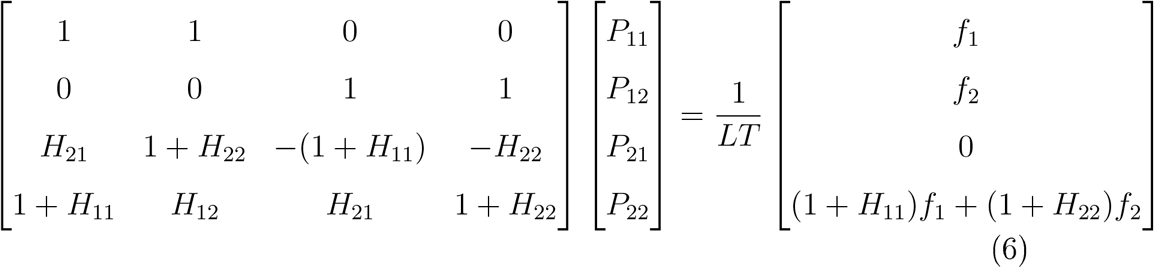

where *L* is the sample width (approximately 20 mm, measured before testing), *T* is the sample thickness in the reference configuration (measured from ultrasound images), and *f*_1_ and *f*_2_ are the forces measured by the BioTester load cells parallel to the *X* and *Y* directions, respectively.

## 3. Results

Volume ultrasound images showed a uniform speckle pattern throughout the tissue volume, with no artifacts from the fishhooks (Fig. 2A). The approximately equibiaxial stretch is apparent in lateral *u*_*X*_ (Fig. 2B) and elevational *u*_*Y*_ displacements (Fig. 2D). The axial displacement (Fig. 2C) exhibits a gradient with depth, indicating thinning of the specimen during stretch.

**Figure 1:**
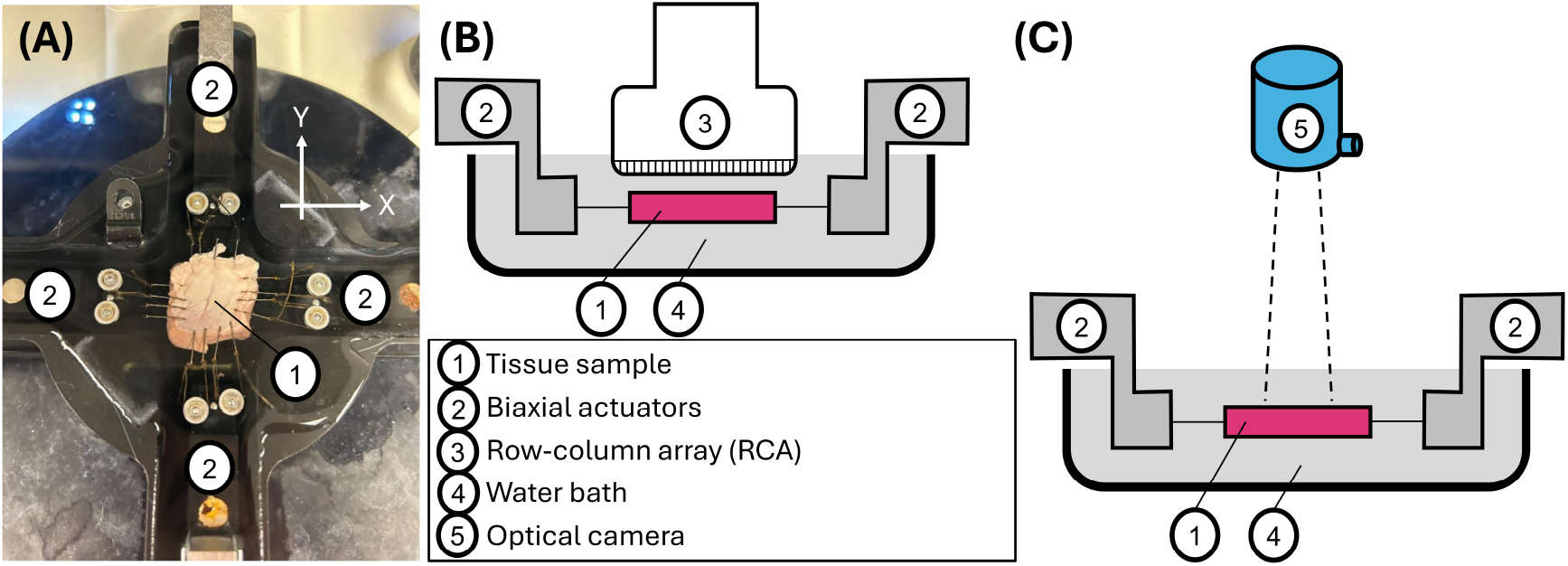
(A) Top-view photograph of excised square sample of RV myocardium loaded into biaxial testing device. Schematics of measurement configurations employing (B) 3D ultrasound and (C) optical camera imaging.

**Figure 2:**
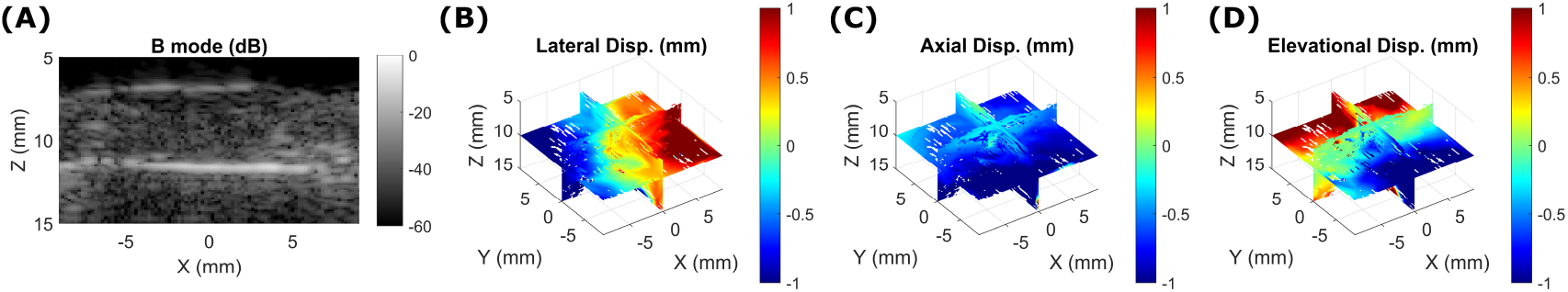
(A) X-Z cross section of the 3D ultrasound image of the LV specimen. (B)-(D) Triplane views of displacement vector components throughout the specimen volume in the maximum stretch for 1-to-1 aspect ratio protocol. Lateral, axial, and elevational displacement refer to displacement in the *X, Z*, and *Y* directions, respectively.

In-plane strain measured by ultrasound is in good agreement with ground truth optical camera-derived strain (Fig. 3A). 3DUST also enables measurement of out-of-plane strain with low uncertainty (Fig. 3B). The mean error of all 36 in-plane strain measurements is 0.7% strain with standard deviation of l.4% strain (Fig. 3C). The root-mean-square error (RMSE) is l.6% strain, similar to our previous study.^8^ The statistically significant error bias (*p* = 0.005) possibly resulted from tissue damage sustained during the first test cycles such that the tissue stretched less for the same actuator displacements during the repeated cycles that were observed by camera.

**Figure 3:**
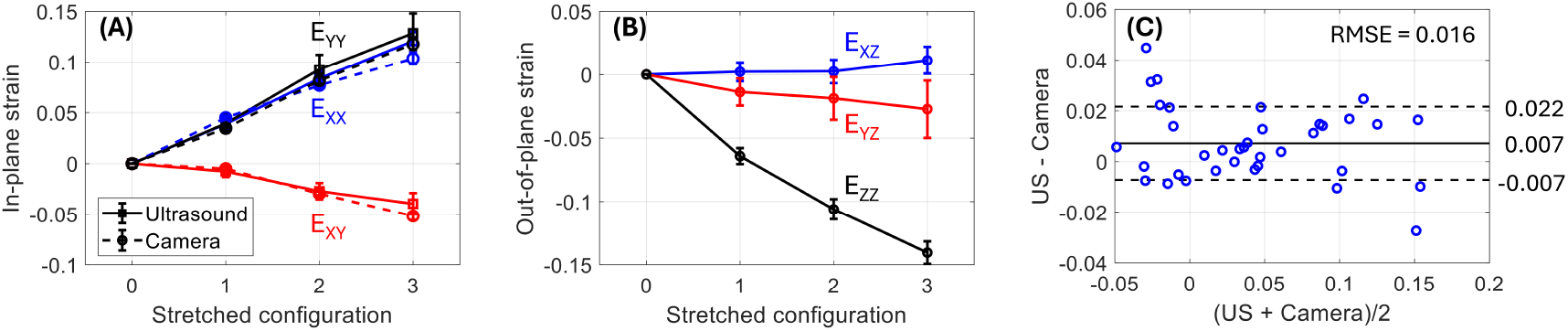
(A) In-plane and (B) out-of-plane strain components computed from 3D ultrasound (solid curves) and optical camera (dashed curves, ground truth) for the LV specimen in 1:1 stretching protocol. Curves in (A) and (B) are straight lines meant to guide the eye between measured data points. (C) Bland-Altman representation of all in-plane strain measurements (blue circles); solid and dashed horizontal lines are mean difference and +/-one standard deviation of difference. Root-mean-square error (RMSE) is 1.6% strain.

**Figure 4:**
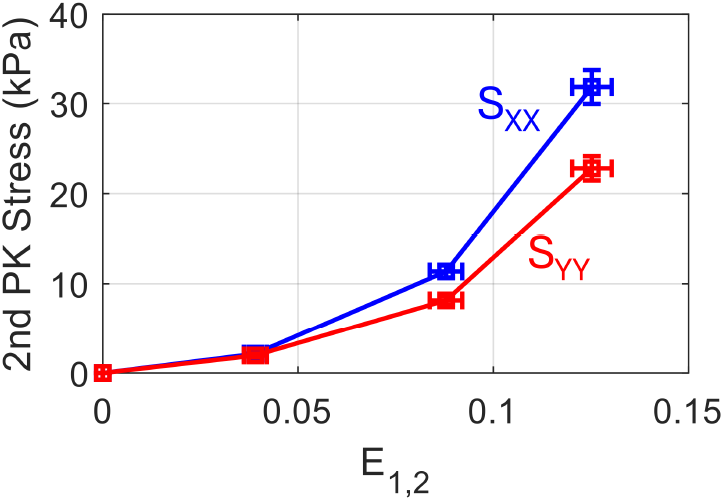
Second Piola-Kirchhoff stresses *S*_*XX*_ and *S*_*Y Y*_ versus mean positive principal strain *E*_1,2_ from 3DUST for the LV specimen in 1:1 stretch protocol. Curves are straight lines meant to guide the eye between measured data points.

Measured stress-strain curves reveal similar stiffness, nonlinearity, and anisotropy as in previous studies of porcine ventricular myocardium.^8,13^

## 4. Discussion

Ultrasound volume imaging with an RCA enables faster 3D imaging than past approaches that used a mechanical scan of a linear array transducer.^8,9^ Faster volume image acquisition can enable more robust biomechanical testing by allowing for observation of a larger number of deformed configurations in a reasonable testing duration. High-frame-rate volume imaging (up to l00 Hz for our sequence) can be used to study dynamic deformations and myocardium viscoelasticity, which are of recent interest.^12^

The major limitation of the approach is processing time. 3DUST calculation for one deformed configuration required several hours, compared to several seconds for computation of surface strain from camera images. Faster cross correlation algorithms can improve 3DUST computation times.^14^

## 5. Conclusion

We implemented a technique for measurement of 3D tissue deformation and six-component tissue strain using ultrasound volume imaging with a row-column array during biaxial mechanical testing of porcine ventricular myocardium. The method was validated against standard ground truth optical camera imaging of the surface strain. The method can improve biomechanical testing of thick myocardial specimens where assumptions of plane stress or incompressibility may not apply.

## Acknowledgments

This work was supported by the Pittsburgh Undergraduate Research Diversity Program (PURDiP) at the University of Pittsburgh Vascular Medicine Institute (NIH grant no. 5R25HL130600), and by the University of Pittsburgh Center for Research Computing.

## Data availability

All data is available from the authors upon reasonable request.

